# LLPSight: enhancing prediction of LLPS-driving proteins using machine learning and protein Language Models

**DOI:** 10.64898/2026.03.01.708814

**Authors:** Valentin Gonay, Rosario Vitale, Georgina Stegmayer, Michael P. Dunne, Andrey V. Kajava

## Abstract

In eukaryotic cells, essential functions are often confined within organelles enclosed by lipid membranes. Increasing evidence, however, highlights the role of membrane-less organelles (MLOs), formed through liquid-liquid phase separation (LLPS). MLO assemblies are typically initiated by “driver” proteins, which form a scaffold to recruit additional “client” molecules. By leveraging expanding MLO datasets and modern machine learning approaches, we developed LLPSight, an ML-based predictor of LLPS-driving proteins. The model was trained using rigorously curated datasets: a positive set of proteins experimentally confirmed to drive LLPS in vivo and a negative set of soluble, unstructured proteins not associated with LLPS. For the features, we employed a cutting-edge approach using embeddings from protein Language Models. LLPSight achieves the highest F1 score (0.885) among existing tools, enabling more efficient discovery of new LLPS drivers eagerly awaited by researchers for experimental validation. An additional key feature of LLPSight is its ability to perform proteome-wide analyses; application to the human proteome yielded promising targets. LLPSight can be obtained from authors upon request.

## Introduction

Liquid-liquid phase separation (LLPS) in cells is a process by which certain proteins and nucleic acids spontaneously separate from the surrounding cytoplasm or nucleoplasm to form dynamic, membrane-less compartments, also called biomolecular condensates or liquid droplets (Fig.1a)^1–5^. These liquid droplets, known as membrane-less organelles (MLOs), perform a variety of critical cellular functions^1^. Among the well-known functional MLOs are stress granules, which form in response to cellular stress (such as oxidative stress, heat shock, or UV radiation). They temporarily sequester mRNAs and several DNA/RNA-binding proteins, pausing transcription and translation to help the cell conserve resources and then rapidly recover from stress^2^. The other known MLOs are P granules, RNA-protein condensates primarily composed of RNA helicases like DDX4/VASA, PGL, and MEG proteins, RNA-binding proteins, and RNAs, functioning in germ cell development and mRNA regulation^3^. MLOs can also be composed purely of proteins, such as Tau-driven LLPS assemblies that concentrate tubulin into condensates to nucleate microtubule bundles^4,5^. The number of such MLO examples is steadily increasing, reflecting the growing interest of biologists in this emerging field of research^6,7^. The growing interest is also justified by evidence that some MLOs can become pathogenic when dysregulated (through mutations, abnormal post-translational modifications, or environmental stress) leading to the formation of pathological condensates associated with various diseases^8,9^.

**Fig. 1:**
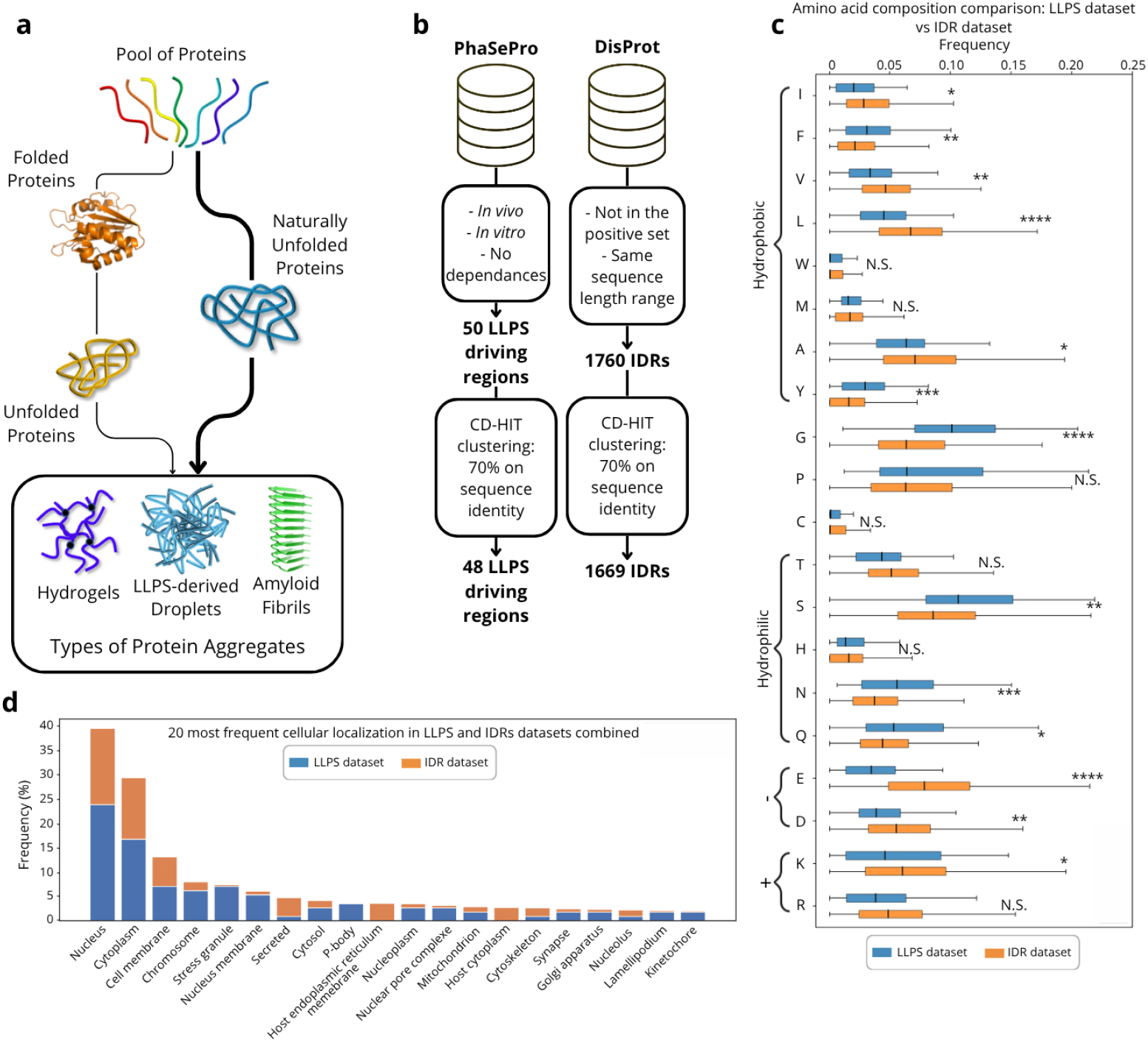
**a**, Simplified scheme of relationship between structural states of proteins and different aggregation types. **b**, Scheme of the construction of the LLPS driver dataset as positive and non-LLPS driving IDR dataset as negative. **c**, Comparison of the amino acid composition of the two datasets (LLPS drivers in blue and above, IDRs in orange and below) (Mann-Whitney U p-value ≥ 0.05: N.S., p-value < 0.05: *, p-value < 0.01: **, p-value < 0.001: *** and p-value < 0.0001: ****). **d**, Comparison of the repartition of the cellular localization among the two datasets (LLPS drivers in blue and below, non-LLPS driving IDRs in orange and above).

Proteins involved in MLOs are commonly classified into two categories. Driver proteins (also called scaffolds or seeds) can undergo LLPS independently^10^, while client molecules, often proteins, but also nucleic acids or small molecules, are recruited into the condensate but do not drive MLO formation on their own^11,12^.

Increasing interest in identifying novel LLPS proteins has led to the development of several computational prediction methods. Such predictors are urgently awaited by the scientific community for their potential to estimate a protein propensity for phase separation, guide experimental validation, and aid in identifying disease-related targets. LLPS predictors generally rely on two major approaches: knowledge-based methods, which use algorithms derived from observed physicochemical properties of proteins involved in LLPS, and more recent machine learning (ML) techniques that classify sequences based on their potential to undergo (or not) phase separation. An example of a knowledge-based LLPS prediction tool is ParSe_v2, which classifies protein regions into three categories: folded, intrinsically disordered, and phase separation-promoting regions^13^. Examples of ML-based methods include PICNIC^14^, FuzDrop^15^, and catGRANULE2.0^16^, which provide binary classification of proteins as LLPS-forming (positive) or non-LLPS-forming (negative).

Training and benchmarking LLPS predictors require datasets of LLPS-related proteins. Some of the most comprehensive sets come from the PhaSePro and DrLLPS databases^17,18^. Despite recent advances, existing predictors require substantial improvement to achieve reliable performance. One promising direction is improving the quality of the training data used to develop these methods. For example, including both client and driver proteins in the positive set, as in PICNIC and catGRANULE 2.0, may reduce the reliability of these tools in predicting the driver proteins most essential for MLO formation. The quality of predictions also depends on the selection of proteins for the negative set. There is considerable flexibility in choosing the negative set, as many proteins do not undergo LLPS or interact with LLPS-forming proteins.

Another promising direction is enhancing the performance of machine learning predictors through more effective feature selection by using the recently proposed protein Language Models (pLMs). pLMs have emerged over the past five years^19–21^, exploiting large amounts of unannotated protein sequence data. These models capture aspects of the “grammar” inherent in protein sequences^22^ and have become a powerful representation method for advancing protein prediction and annotation^23^. For many applications, alignment-free pLM-based predictions now achieve significantly higher accuracy than classical models. A pre-trained pLM can convert a raw protein sequence into a dense feature vector (embedding) that encodes its representation. This embedding can serve as the sole input for downstream supervised models, simplifying modeling and enabling predictive models to learn specific target tasks directly from the embeddings.

In this study, we developed a new ML-based LLPS predictor that leverages pLM embeddings, which was trained specifically on rigorously selected LLPS-driving regions of proteins known to undergo phase separation *in vivo* and *in vitro*, aiming for greater accuracy within this protein category. Because LLPS drivers are predominantly intrinsically disordered, we chose a negative set composed of intrinsically disordered regions and proteins (IDRs/IDPs) that do not undergo LLPS^24^ (Fig.1a). Unlike other predictors, this design explicitly contrasts disordered LLPS drivers with disordered non-LLPS proteins.

## Results

### Construction of Datasets

Our positive dataset of LLPS-driving protein regions was collected from the PhaSePro database^17^. We have chosen PhaSePro as our source because of its high level of completeness, detailed annotation, and the ease of selecting only LLPS drivers through its searchable interface. To build the LLPS-driver set, we extracted all entries reported to undergo LLPS independently, i.e., without requiring additional partners. We extracted 50 entries, which were clustered at 70% sequence identity to reduce redundancy, yielding a final set of 48 entries (Fig.1b).

The negative set was composed of experimentally validated soluble IDRs and IDPs retrieved from the DisProt database^25^. This choice is justified by the fact that LLPS-driving regions of proteins are overwhelmingly disordered^24^. Consequently, using proteins with stable structures as the negative set risks training the model to predict disordered regions rather than true LLPS-driving regions. At the same time, unstructured LLPS drivers exhibit distinct sequence features compared with soluble IDRs; therefore, using IDRs as the negative set is the most appropriate approach for training and benchmarking ML methods.

We ensured that none of the entries in the negative set overlapped with those in the positive dataset. Negative entry sequences were selected to match the length distribution of the positive set (35-958 vs. 35-995 residues). After clustering at 70% sequence identity, the negative set comprised 1,669 entries (Fig.1b). Both sets are primarily from *Homo sapiens* and *Saccharomyces cerevisiae*, with other species in smaller numbers that present significant differences in amino acid composition and protein cellular localization (Fig.1c,d).

### Training/Testing set split

Because the negative set contained much more entries than the positive set, it was randomly sub-sampled to create a class-balanced dataset, maintaining a 1:1 ratio between positive and negative entries (Fig.2a). To determine the required training set size, we computed a learning curve using the Scikit-learn Python library (version 1.3.1) with a raw (non-optimized) ExtraTrees classifier and monitored performance with the F1 score (Fig.2c). The learning curve allowed us to identify the training set size at which performance reached a plateau, while still retaining enough entries for testing. A 70/30 split between training and testing sets generally yields representative results. Accordingly, from a total of 96 entries (48 positive and 48 negative), we selected 66 entries (33 positive and 33 negative) for training, corresponding to the plateau point, and reserved 30 entries (15 positive and 15 negative) for testing (Fig.2a).

**Fig. 2:**
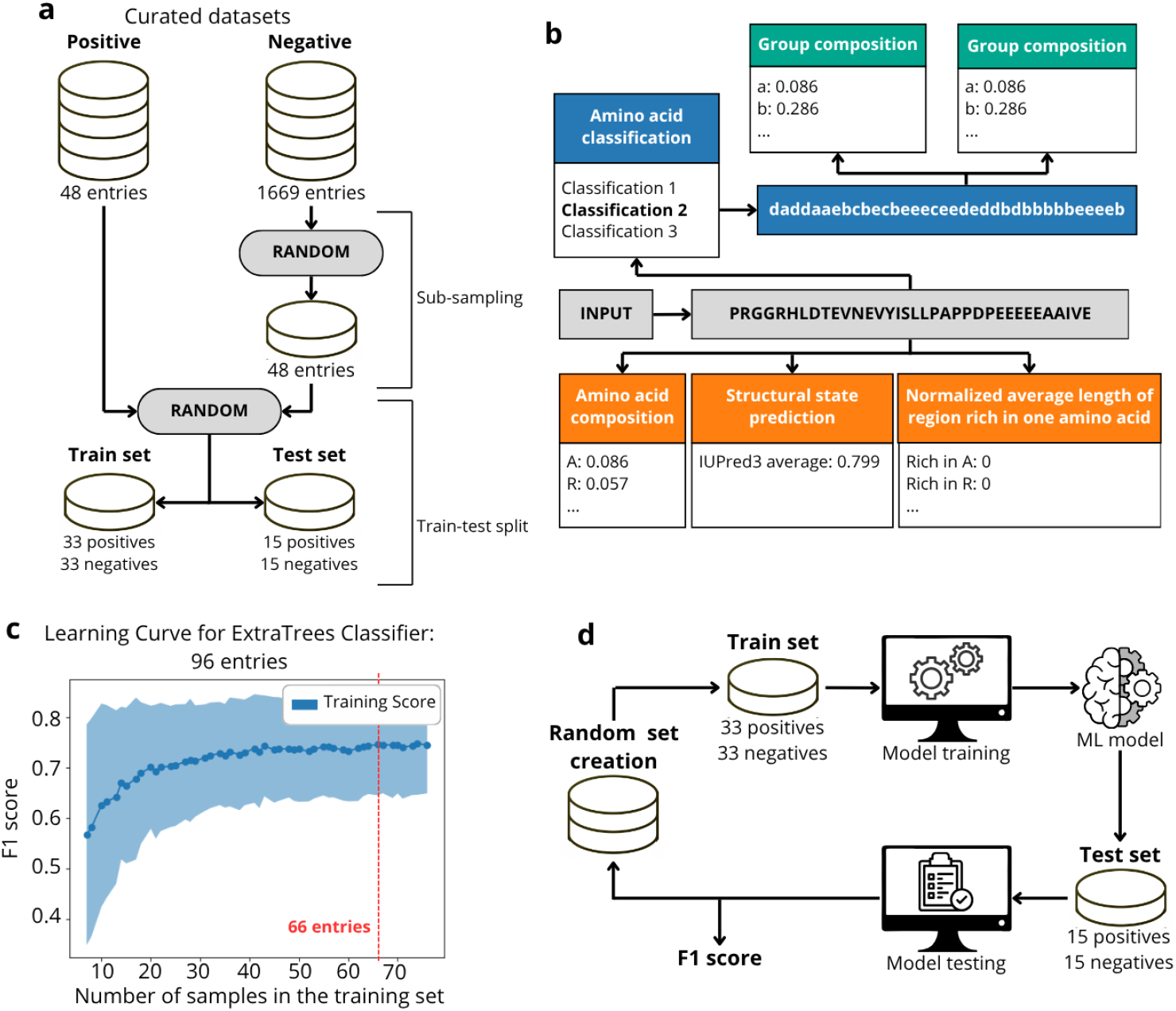
**a**, Flowchart of the generation of the training and testing sets from the positive and negative data obtained after clustering. **b**, Schematic representation of generated knowledge-based features from an input sequence. Two classes of features are represented in orange (amino acid composition, structural state prediction, and normalized average length of region rich in one type of amino acid) and green (Group composition and group transition). The first class is generated from an amino acid sequence, while the second one is derived from the sequence generated by amino acid classification. **c**, Learning curve showing the evolution of the obtained F1 score depending on the number of samples used in the training set. Dots represents the average F1 score for a 200-fold cross-validation from the in-built option in the scikit-learn library. Standard deviation is represented as the blue area above and under the dots. **d**, Schematic representation of the cross-validation process. This process is repeated as much as needed starting at the random set generation (corresponding to the **Figure 2a**) and all F1 scores are stored. The average from the stored F1 scores is then used as a final score.

### Feature Selection and pLM embeddings

As an initial approach to the prediction problem, and given the strong performance of earlier ML methods using knowledge-based features for identifying amyloid-forming regions^26^, we adopted a similar strategy (Fig.2b, Table 1). Because the ML model receives amino acid sequences as input, we focused on sequence-based features. We used 20 features representing individual amino acid frequencies and 5 features based on frequencies of amino acid groups with similar physicochemical properties. Comparing model performance across three group classifications (Supplementary Table 1), we selected the most effective: group a (R, H, K), group b (D, E), group c (S, T, N, Q), group d (C, G, P), and group e (A, V, I, L, M, F, Y, W). We then derived 25 group-transition features capturing pairwise transitions between adjacent groups (5 × 5 = 25 combinations).

**Table 1:** Definitive set of knowledge-based features. A set of features selected after a statistical comparison of value distribution between the positive and the negative sets

| Feature type | Feature |
| --- | --- |
| Structurality (1) | IUPred score |
| Amino acid composition (12 of 20) | 'N', 'D', 'C', 'Q', 'E', 'G', 'H', 'F', 'P', 'S', 'W', 'Y' |
| Amino acid group composition (4 of 17*) | 'grp_A', 'grp_B', 'grp_C', 'grp_D' |
| Amino acid group transition (11 of 25) | 'A_to_D', 'B_to_E', 'C_to_C', 'C_to_D', 'C_to_E', 'D_to_A', 'D_to_C', 'D_to_D', 'D_to_E', 'E_to_B', 'E_to_D' |
| Normalized Average length of regions rich in one amino acid (13 of 20) | 'length_RichRegion_Y', 'length_RichRegion_R', 'length_RichRegion_G', 'length_RichRegion_Q', 'length_RichRegion_S', 'length_RichRegion_F', 'length_RichRegion_C', 'length_RichRegion_M', 'length_RichRegion_T', 'length_RichRegion_P', 'length_RichRegion_H', 'length_RichRegion_E', 'length_RichRegion_N' |

\* The total number of groups includes three amino acid classifications (Supplementary Table 1)

Because LLPS-driving regions are often enriched in specific residues^2^, we generated features to capture this property. Residue-frequency differences were computed within sliding windows of varying sizes between the positive or negative sets and all IDRs from DisProt^25^, allowing determination of the optimal window and enrichment threshold. Using a 10-residue window, a region was considered enriched when the frequency of a residue in positive or negative sequences was at least twice that in DisProt. This binary scheme yielded 20 features.

As LLPS typically arises from IDRs, we also included the average disorder score of each sequence^27^. In total, 71 features were derived for subsequent selection of the optimal combination to classify LLPS-driving proteins versus soluble IDRs. Only features significantly separating the positive and negative sets were retained, resulting in 41 high-impact knowledge-based features for model development (Table 1).

As an alternative to the knowledge-based features model, we also developed another model based on the more recent promising approach for ML models: the use of embeddings from pLMs^23^. To generate embeddings, we followed the protocol of Vitale et al.^28^ and selected the two top-performing pLMs for protein classification: ESM and ProtTrans according to Fenoy et al.^29^. The ESM family of models was pre-trained using amino acid sequences as inputs^30^. The most recent model, ESM2^31^, is a general-purpose model with up to 15 billion parameters, making it the largest pLM to date. We used esm2_t33_650M_UR50D, which generates embeddings of size 1,280 for each protein sequence. ProtTrans ProtT5^32^ is an encoder-decoder model that projects a source language into an embedding space and then generates a translation to a target language. Among several ProtTrans models, we used ProtT5-XL-U50 (Rostlab/prot_t5_xl_half_uniref50-enc), indicated as the best by the authors, which provides embeddings of size 1,024 per protein sequence.

Next, we compared model performance using either knowledge-based features or pLM embeddings.

### Model selection

To identify the best ML predictor, we evaluated six supervised classifiers: AdaBoost, DecisionTree, ExtraTrees, GradientBoosting, RandomForest, and Support Vector Machine (SVM). Each model was optimized using a random grid search over 1,000 hyperparameter combinations, with average performance assessed across 75 cross-validation runs. The hyperparameter set yielding the highest F1 score was retained for model selection. For final model selection, each classifier was evaluated with 500 cross-validation runs using the optimized hyperparameters. The resulting F1 score distributions were statistically compared to identify the best-performing model.

Overall, embeddings achieved higher performance than knowledge-based features (Fig.3a). Among pLM embeddings, ESM2 outperformed ProtT5 by a small margin. The average performance of each optimized model with its best feature set was as follows: Adaboost with ESM2 (F1 score = 0.872), DecisionTree with ESM2 (F1 score = 0.746), ExtraTrees with ESM2 (F1 score = 0.886), GradientBoosting with ESM2 (F1 score = 0.877), RandomForest with ESM2 (F1 score = 0.885) and SVM with ProtT5 (F1 score = 0.883) . Based on these results, the top-performing models with no significant differences in F1 score were Random Forest (ESM2), ExtraTrees (ESM2), Gradient Boosting (ESM2), and SVM (ProtT5). Random Forest and ExtraTrees achieved the highest scores. We selected Random Forest with ESM2 to build our predictor, LLPSight, since this type of model can reliably identify optimal split points (thresholds) for each feature.

**Fig. 3:**
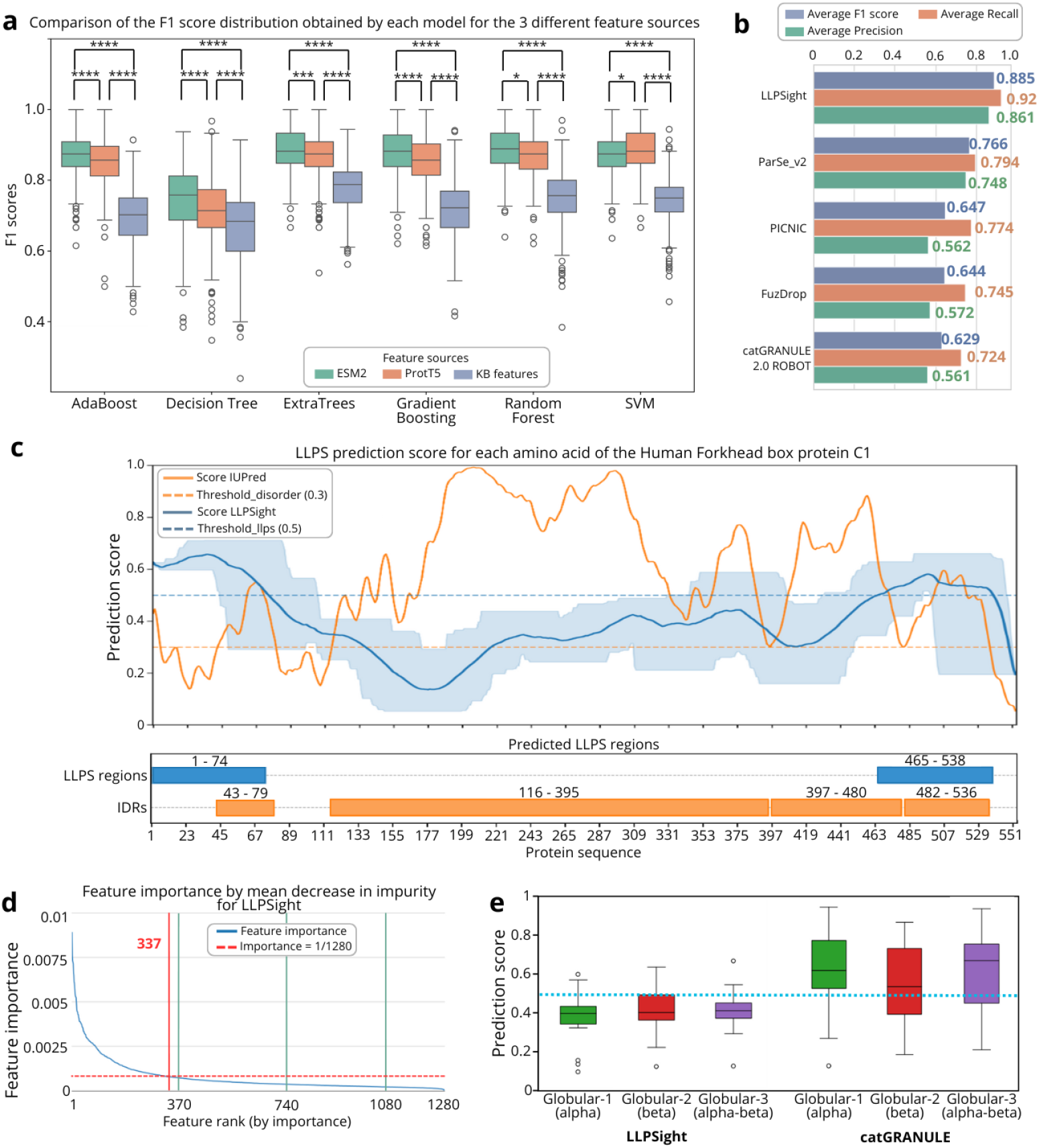
**a**, F1 score distribution obtained after 500 cross-validation sessions on optimized models for three different sources of features: ESM2 (green, left), ProtT5 (orange, middle) and our knowledge based (KB) features (blue, right). All different sources gives significantly different results (Mann-Whitney U p-value ≥ 0.05: N.S., p-value < 0.05: *, p-value < 0.01: **, p-value < 0.001: *** and p-value < 0.0001: ****). Statistic comparison of each best source of features for every model available in Supplementary Table 2. **b**, Average result obtained by the benchmarked models (average based on 10 groups randomly composed of 17 positive and 17 negative entries) and average performance of LLPSight after 500 sessions of cross-validation. Three metrics are displayed: the average F1 score (blue, top), the average recall (red, middle), and the average precision (green, bottom). Detailed scores obtained for each set are available in Supplementary Table 4. **c**, Prediction result for the Human Forkhead box protein C1 (UniProt: Q12948). The upper box is a graphical representation of the score distribution along the sequence for IUPred3 score (orange) and for LLPSight score (blue) with the average scores as a solid line surrounded by the maximum and minimum. The lower box represents the LLPS driving regions (blue, top), IDRs (orange, bottom). LLPS driving regions are defined as at least consecutive 12 residues with a score (defined by the user, can use the maximal, average (default) or minimal) higher than the threshold (0.5 by default). **d**, Features ranking by mean decrease in impurity for LLPSight. The red mark at 337 shows the number of features with an importance higher than 1/1280. Green markers at 370, 740, and 1080 correspond to 10× the number of training positives, training samples, and total samples, respectively. **e**, Prediction score distributions of LLPSight (right) and catGRANULE (left) for three groups of globular structures obtained from the CATH database. The blue dotted line at 0.5 represents the threshold for the classification.

Feature importance was analyzed to assess potential overfitting due to the high-dimensional embeddings. Using the mean decrease in impurity (Scikit-learn v1.3.1), 337 of 1,280 features exceeded the threshold of 1/1,280 (Fig.3d), indicating that the remaining embeddings contribute minimally and are unlikely to drive overfitting.

### Benchmark results

We evaluated LLPSight performance against previously published LLPS predictors, ParSe_v2, catGRANULE 2.0 ROBOT, FuzDrop, and PICNIC, using the benchmark set (Supplementary Table 3). LLPSight demonstrated superior performance, as reflected by highest F1 score (0.89), recall (0.92), and precision values (0.86) (Supplementary Table 4, Fig.3b). Recall is high across all methods (0.72-0.92), showing they effectively identify LLPS-driving proteins. Our method stands out in precision, reducing false positives from non-LLPS proteins (soluble IDRs/IDPs) compared with other approaches.

To evaluate whether our method can distinguish LLPS drivers from globular proteins, we extracted PDB identifiers from the CATH database^33^ for proteins primarily composed of α-helices (Globular-1), β-strands (Globular-2), or a mixture of both (Globular-3). Thirty proteins from each group were randomly selected, and their LLPS propensity was predicted. LLPSight classified only 10%, 26.7%, and 6.7% of Globular-1, -2, and -3 proteins as positive (Fig.3e), confirming its discriminative ability. In contrast, the other method, catGRANULE, which is suitable for analyzing large numbers of sequences, assigned high LLPS scores to many globular proteins (Globular-1: 83%, Globular-2: 53%, Globular-3: 66%) (Fig.3e).

### Additional functionalities of LLPSight

#### Sliding Window for Prediction of LLPS-driving regions

Although LLPSight is trained on LLPS-driving regions, its goal is to analyze full-length proteins. We therefore adapted the model to classify regions within long sequences as LLPS-driving or not, using a default prediction window of 50 residues. The window is set to 50 residues (rounded from the shortest region in the training set, which was 35 residues) to efficiently detect relatively short regions in proteins without excessive computation. This window size can be modified by the user. The window slides from the N-to the C-terminus in steps of one residue. LLPSight calculates a prediction score for each window, and each amino acid residue therefore receives multiple scores, one from each window that covers it. The final output reports the maximum, average, and minimum score per residue (Fig.3c). A protein is classified as an LLPS driver if at least one region containing 12 or more consecutive residues has an average score above 0.5.

In many cases, LLPS-driving regions overlap with IDRs. However, in general, the regions predicted to drive LLPS do not perfectly match the IDRs predicted by IUPred3. To better illustrate this difference, we output both the LLPS-predicted regions and the IUPred3-predicted IDRs and provide a graphical display, allowing a clear visual comparison of the two types of predictions (Fig.3c).

#### Transmembrane helices

It was observed that LLPS-driving regions tend to be of low complexity and enriched in a few specific amino acids. At the same time, low-complexity sequences enriched in hydrophobic residues can form transmembrane helices rather than drive LLPS. This subtle distinction is often not captured by current LLPS prediction methods. To alert users, we highlight potential transmembrane helices identified using the simplified filter described by Falgarone et al.^34^ (Fig.3c).

### Human proteome analysis by LLPSight

An important feature of LLPSight is its ability to perform proteome-wide analyses. Among the predictors tested (Fig.3b), only catGRANULE 2.0 ROBOT^16^ offers comparable functionality. Both tools were applied with default parameters to analyze the human proteome downloaded from UniProt using the canonical sequences option. LLPSight predicted 1598 (7.9%) LLPS drivers among human proteins (Fig.4a). In comparison, catGRANULE 2.0 ROBOT^16^ predicted over half of the human proteome (52.2%) to be involved in LLPS formation (Fig.4b). Experimental evidence to date indicates that, although LLPS driving organelles play important biological roles, they remain a relatively rare phenomenon. This suggests that catGRANULE may substantially overpredict LLPS drivers, as it is unlikely that every second human protein contributes to LLPS.

**Fig. 4:**
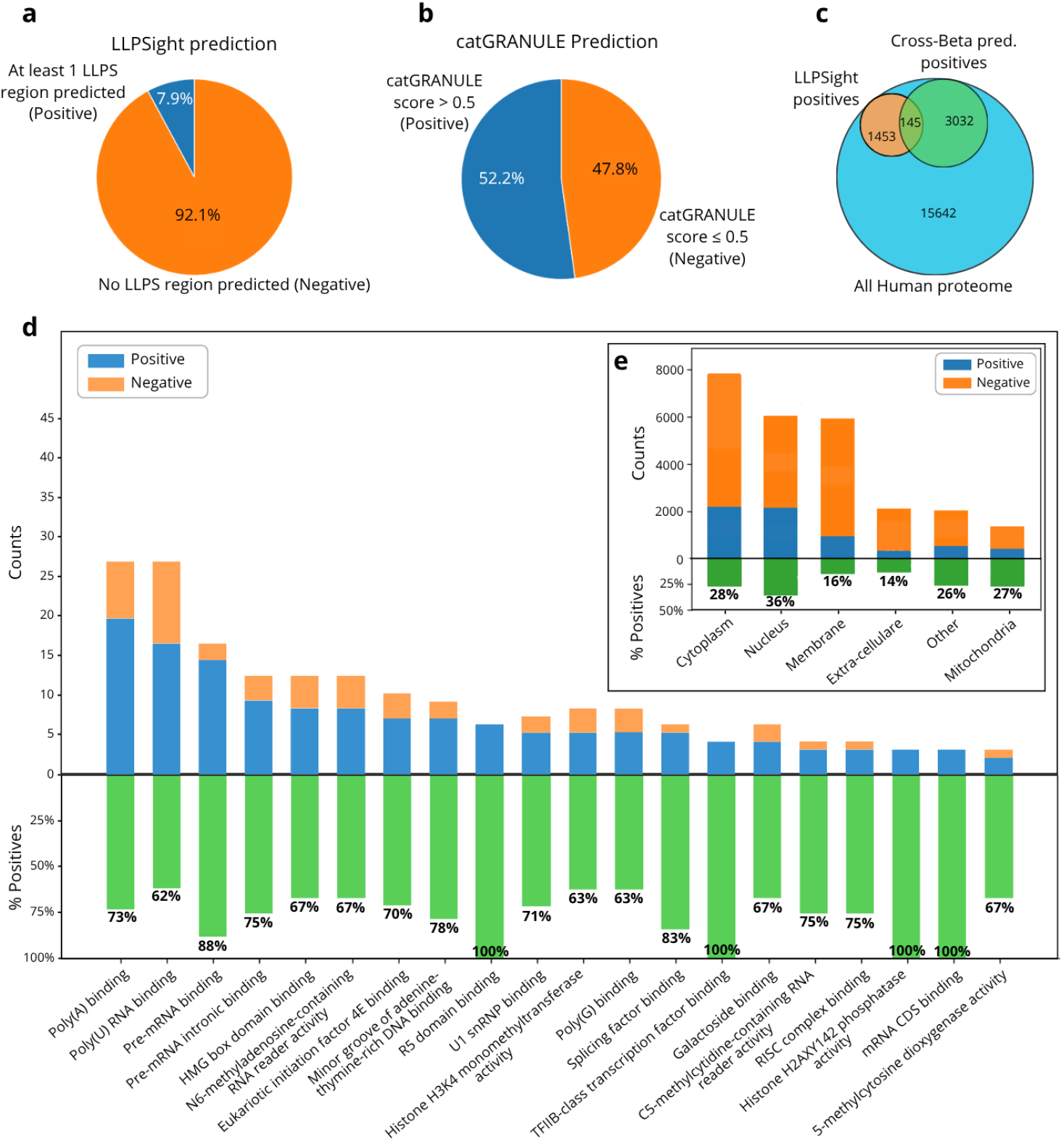
**a**, Proportion of positive (blue) and negative (orange) prediction from the LLPSight results on the human proteome. The definition of positive is set to all sequences presenting at least one predicted LLPS prone region. **b**, Prediction from the catGRANULE 2.0 ROBOT results on the human proteome. The definition of positive prediction corresponds to sequences with a score higher than 0.5. **c**, Venn diagram representing the overlap between the positive prediction from LLPSight with at least one LLPS region predicted (orange), the positive prediction from the Cross-Beta predictor of amyloidogenicity (green) and the entire human proteome (blue). **d**, Comparison of the positive (blue, bottom) and negative (orange, top) protein functions based on ontologies for functions where at least 60% of proteins are predicted positive. The proportion of positive protein by function is expressed in percentage and shown in green. **e**, Comparison of the subcellular locations of the positive and negative prediction groups in six major locations: cytoplasm, nucleus, membrane (all kinds of membrane), extra-cellular, mitochondria, and others. The count comparison shows the number of each class (blue and bottom for positive, orange and top for negative) while the proportion of positive is displayed in percentage (green).

The 1598 potential LLPS-driving proteins predicted by LLPSight were analyzed in terms of their amino acid composition, subcellular localization, and functions. LLPS driving regions showed significant enrichment of Gly, Pro, Ser, and Gln, with other residues being underrepresented relative to non-LLPS regions. The amino acid composition of LLPS-driving regions differs significantly from ones of non-LLPS and IDRs Supplementary Figure 1-2 The proportion of residues belonging to LLPS regions occupy, on average, 1.6% of the total proteome. Subcellular localization from UniProt was grouped into six categories: Nucleus, Cytoplasm, Membrane, Extracellular, Mitochondria, and Other. As shown in Fig.4e, LLPS proteins are most frequently localized in the nucleus (36% of nuclear proteins), whereas membrane proteins show the lowest representation (14%). This is consistent with the idea that membrane proteins, particularly those with transmembrane domains, are unlikely to contain LLPS-driving regions, as their localization restricts involvement in membraneless organelles.

Functional associations of LLPS drivers were obtained from Gene Ontology (GO) annotations^35^. Functions in which at least 60% of proteins were predicted as LLPS-driving were predominantly related to RNA binding, consistent with the frequent formation of protein-RNA membraneless organelles, such as stress granules (Fig.4d).

We also compared our LLPSight predictions with those obtained from the Cross-Beta predictor^26^, which detects proteins capable of forming amyloids under physiological conditions. The results show that approximately 10% of LLPS drivers are also capable of forming amyloid fibrils (Fig.4c), highlighting the overlapping sequence properties of amyloid- and LLPS-forming proteins.

To identify novel LLPS drivers predicted exclusively by LLPSight, we focused on proteins absent from positive LLPS datasets (PhaSePro^17^, DrLLPS^18^), our negative dataset from DisProt^25^, and not predicted as positive by catGRANULE. This analysis identified 528 novel LLPS drivers, (each containing at least one LLPS-prone region) that could be further investigated experimentally. Refinement of this protein set can be achieved by assuming that functionally important LLPS-driving regions are conserved in orthologs from related species.

For example, the newly identified LLPS driver Decreased Expression in Renal and Prostate Cancer protein (DERPC) was predicted to have LLPS propensity in multiple species, including *Homo sapiens, Mus musculus, Rattus norvegicus, Bos taurus, Vulpes vulpes, Phyllostomus discolor, Mustela putorius furo*, and *Mesocricetus auratus*, suggesting that its LLPS-driving region is functionally conserved. This protein may represent a promising target for experimental investigation. In contrast, although LLPS propensity was predicted for Spermatogenesis-associated protein 3 (SPATA3) from *Homo sapiens*, orthologs from closely related species (*Daubentonia madagascariensis, Ceratotherium simum simum, Physeter macrocephalus, Sousa chinensis, Panthera pardus, Puma concolor, Castor canadensis, Cricetulus griseus, Dipodomys ordii, Tursiops truncatus*, and *Neophocaena asiaeorientalis asiaeorientalis)* lacked LLPS-driving regions. This protein may be a less promising target.

## Discussion

With the advent of ML-based computational tools such as AlphaFold^36^, protein structural annotation based solely on sequence analysis has improved dramatically. These predictions are particularly reliable for proteins that fold into stable 3D structures. However, in recent years, scientists have turned their attention to a different and highly dynamic mode of protein organization: proteins forming membraneless organelles (MLOs), where they remain together within droplets through transient, weak interactions. In this “liquid-like” state, the proteins are not fixed relative to one another. Such dynamic arrangements cannot be predicted by structure-focused tools like AlphaFold, highlighting the need for new sequence-based methods to identify these proteins.

As LLPS is receiving increasing attention, such predictors are urgently needed by the scientific community to evaluate our understanding of LLPS and its prevalence in living organisms, estimate the frequency of proteins with LLPS propensity, identify novel targets for experimental validation, and facilitate the study and prediction of missense mutations in LLPS-driving regions associated with hereditary diseases.

Current predictors of LLPS-related proteins require enhancement. Here, we developed a novel ML-based method, LLPSight, which outperformed existing approaches. Our strategy relied on three key cornerstones: (i) focusing on LLPS drivers, the key proteins that trigger MLO formation; (ii) including in the negative set IDRs that are known to be soluble and lack evidence of participating in LLPS; and (iii) exploring a wide range of approaches to generate ML features, including embeddings from pLMs.

An additional filter was applied to the positive dataset, to ensure that LLPS-driving entries have both *in vitro* and *in vivo* experimental evidence of self-assembling LLPS droplets. Features for ML models were generated using either knowledge-based approaches or embeddings from pMLs. Six supervised models were tested, and those based on ESM2 embeddings outperformed the knowledge-based features. Consequently, a Random Forest model with ESM2 embeddings was selected to build LLPSight, which is, to our knowledge, the only LLPS predictor using pLM embeddings as input.

Benchmarking LLPSight against four other published tools showed that it achieved the highest performance (F1 score 0.89, recall 0.92, and precision 0.86) when distinguishing LLPS-driving proteins from intrinsically disordered, non-LLPS-driving proteins.

Using LLPSight, we performed a large-scale analysis of the human proteome and identified approximately 8% of proteins as potential LLPS drivers. These proteins are characterized by enrichment of Gly, Pro, Ser, and Gln in LLPS-driving regions; they are most frequently localized in the nucleus and least in membranes, and their functions are predominantly related to RNA binding. Importantly, hundreds of newly identified potential LLPS drivers are now available for experimental investigation.

LLPSight can be used as a command-line-based application available from authors upon request. It accepts as input a protein sequence or a FASTA file, including multi-FASTA format. Query sequences must be at least 20 residue long and composed exclusively of natural amino acids. Prediction thresholds can be modified by the users, and results are provided a .csv file with optional graphical display Potential transmembrane regions are highlighted as a warning, since they may prevent the formation of LLPS droplets.

## Methods

### Data Selection and Training/Testing Set Splitting

LLPS-driving protein regions were collected from the PhaSePro database^17^ (downloaded on April 5, 2024) by selecting entries reported without dependencies. Extraction was performed using a Python script, selecting entries where *in_vivo* and *in_vitro* were True, and *partner_dep, rna_dep, ptm_dep*, and *domain_dep* were set to ‘N’. For the negative set, we selected soluble IDRs and IDPs from the DisProt database (downloaded April 8, 2024)^25^. Entries overlapping with the positive set were excluded, and sequences were matched to the positive set length distribution to minimize bias. The initial set of 1,760 entries was reduced to 1,669 after clustering with CD-HIT^37^ at 70% sequence identity. To obtain a class-balanced dataset, the negative set was randomly sub-sampled to achieve a 1:1 ratio with the positive set. Training set size was determined via a learning curve computed with the Scikit-learn Python library (v1.3.1) using a raw ExtraTrees classifier and the F1 score (Fig.2a).

### Features

#### Knowledge-based features

Selection of high-impact knowledge-based features was performed by comparing the statistical distributions of each feature between the positive and negative sets. Only features that showed a significant difference between the two sets were retained for subsequent model development. All statistical analyses have been made in two steps: first, a normality test was made using the Shapiro-Wilk test^38^. If one of the compared distributions was significantly different (p-value < 0.05) from a normal distribution, a Mann-Whitney U rank test^39^ was performed to compare the two distributions. Otherwise, if both distributions were not significantly different from a normal distribution (p-value ≥ 0.05), a Student’s T-test was performed. All tests and statistical analyses were performed using the `SciPy` Python library (version 1.12.0)^40^.

#### pLM embeddings

Embeddings were obtained following Vitale et al.^28^ using the two top-performing pLMs for classification: ESM and ProtTrans^29^. Both models, pre-trained on millions of UniProtKB sequences, generate per-residue embeddings that capture contextual information and hidden dependencies between amino acids^22,23^. These per-residue representations were averaged to produce per-protein embeddings for downstream supervised learning. We used esm2_t33_650M_UR50D, which generates embeddings of size 1,280 per protein sequence, and ProtT5-XL-U50 (Rostlab/prot_t5_xl_half_uniref50-enc), which generates embeddings of size 1,024 per sequence.

### Cross-Validation procedure

To get an average performance not dependent on the used entries in the training and testing sets, we analyze the average F1 score obtained over a representative number of training/testing sessions with, each time, a different random sampling of data present in the training and testing sets. This means that positive data (limited in number) would be split differently between the two sets, and the negative data selected would be different for each session. Every session represents a new model trained and tested with different entries, leading to an average score over those models representing a final score, independent of the choice of data for the sets (Fig.2d). This procedure was used during the next steps of model optimization and selection.

### Model optimization and selection

All models were provided by the Python library `Scikit Learn`^41^. We pre-selected six classifier models: AdaBoost, DecisionTree, ExtraTrees, GradientBoosting, RandomForest, and Support Vector Machine classifier (SVM). These models were first optimized following a random grid search strategy. Each model’s performance has been tested for 1000 random hyperparameter grids, and the average performance was calculated for 75 cross-validation sessions. The combination of hyperparameters showing the highest F1 score has been selected for the model selection process.

For final model selection, each optimized model was evaluated with 500 cross-validation runs using the chosen hyperparameters. Statistical comparisons of the resulting F1 score distributions were performed in two steps: a normality test using the Shapiro-Wilk test, followed by a paired comparison of all model distributions using the Mann-Whitney U rank test if one of the distributions didn’t follow a normal distribution or a Student T-test if both compared distributions were normal.

### Benchmarking

The performance of other LLPS predictors was evaluated using the same positive and negative datasets as for our model. The predictors selected for benchmark were: ParSe_v2^13^, catGRANULE 2.0 ROBOT^16^, FuzDrop^15^, and PICNIC^14^. Those tools have been selected for their respective diversity in algorithms and data used during their development. For instance, ParSe_v2 was developed without the usage of machine learning models and it is based on three sets of data: Folded, Intrinsically Disordered, and Phase-separating as described in Ibrahim et al.^42^. catGRANULE 2.0 ROBOT is a ML-based algorithm using human proteins known to be involved in LLPS as positive examples and proteins not involved in LLPS as negative examples for its training set^16^. FuzDrop has been developed using positive droplet-promoting regions collected from diverse databases (including PhaSePro), and the negative data (non-droplet-forming proteins) were collected from the human proteome available in the Swiss-Prot database. This predictor is also based on a ML approach, which uses a binary logistic regression model^15^. Finally, PICNIC is another recent ML-based binary classifier identifying proteins involved in the formation of condensates and proteins that are not^14^.

To test those tools, we chose to reproduce the conditions used for our testing set: 17 positive entries and 17 negative entries, randomly selected to create 10 cross-validation benchmark groups. All tools were compared using the average F1 score obtained from the test of the 10 groups. FuzDrop only analyzes sequences longer than 45 residues; shorter sequences were treated as negatives (True Negative or False Negative). In addition, FuzDrop produced errors of unknown origin for a few sequences, which were ignored when computing the F1 scores.

## Code availability

LLPSight is open source and can be obtained from authors upon request

## Acknowledgements

This work was supported in part by a CIFRE PhD fellowship through the “Association Nationale de la Recherche et de la Technologie” (ANRT) program, in collaboration between PROTERA and “Centre National de la Recherche Scientifique” (CNRS) to V.G., by EU COST Action ML4NGP CA21160 to G.S., R.V., V.G and A.V.K., by EU Horizon Europe MSCA Staff Exchange project IDPfun2 -grant agreement No. 101182949 to V.G and A.V.K. and by the Fondation pour la Recherche Médicale (grant MND202310017898) to A.V.K. This work was also supported in part by ANPCyT PICT 2022 [grant number #0086] and CAID-UNL 2024 [grant number #0100097] to G.S.

## Author contribution

Conceptualization A.V.K., G.S., M.P.D., V.G.; Methodology A.V.K., G.S., M.P.D., V.G.; Software V.G.; Validation R.V.,V.G.; Formal analysis V.G.; Data curation V.G.; Writing – original draft A.V.K., V.G.; Writing – review & editing G.S., M.P.D.; Visualization A.V.K., V.G.; Supervision A.V.K., G.S., M.P.D.; Project administration A.V.K.; Funding acquisition A.V.K., G.S., V.G.

## Competing interests

The authors declare no conflicts of interest.

## Notes

### Competing Interest Statement

The authors have declared no competing interest.

