## Supplemental Data for "LLPSight: enhancing prediction of LLPS-driving proteins using machine learning and protein Language Models"

**SUPPLEMENTARY DATA. Gonay et al., 2026**

**Supplementary Table 1:** Classification table for amino acids based on their physico-chemical properties. Description of the 3 different classification methods used.

| Classification method | Group name | Classified amino acids |
|---|---|---|
| method 1 | grp_A | R, H, K, D, E |
| | grp_B | S, T, N, Q |
| | grp_C | C, G, P |
| | grp_D | A, V, I, L, M, F, Y, W |
| method 2 | grp_A | R, H, K |
| | grp_B | D, E |
| | grp_C | S, T, N, Q |
| | grp_D | C, G, P |
| | grp_E | A, V, I, L, M, F, Y, W |
| method 3 | grp_A | N, Q |
| | grp_B | A, I, M, V, L |
| | grp_C | C |
| | grp_D | S, T, H |
| | grp_E | D, E, K, R |
| | grp_F | Y, F, W |
| | grp_P | P |
| | grp_G | G |

**Supplementary Table 2:** Statistic result for the comparison of F1 scores obtained by the 6 different optimized models for 500 cross validation sessions.

| Models | Adaboost (ESM2) | Decision Tree (ESM2) | ExtraTrees (ESM2) | Gradient Boosting (ESM2) | Random Forest (ESM2) | SVM (ProtT5) |
|---|---|---|---|---|---|---|
| Adaboost (ESM2) | | | | | | |
| Decision Tree (ESM2) | $1.827 \times 10^{-92}$ | | | | | |
| ExtraTrees (ESM2) | $2.387 \times 10^{-4}$ | $5.751 \times 10^{-108}$ | | | | |
| Gradient Boosting (ESM2) | $0.130$ | $9.645 \times 10^{-99}$ | $0.029$ | | | |
| Random Forest (ESM2) | $6.088 \times 10^{-4}$ | $2.980 \times 10^{-107}$ | $0.808$ | $0.056$ | | |
| SVM (ProtT5) | $3.686 \times 10^{-3}$ | $5.765 \times 10^{-8}$ | $0.410$ | $0.166$ | $0.574$ | |

**<u>Supplementary Table 3:</u>** Benchmark set and sub groups used to evaluate and compare the performances of the different LLPS predictors

SEE CSV “Supplementary Tables.csv”

**Supplementary Table 4:** Details of the obtained scores for each set by each tested LLPS predictors. (**A**) The recall obtained for each set by each predictors. (**B**) The precision obtained for each set by each predictors. (**C**) The F1 score obtained for each set by each predictors.

| (A) | ParSe_V2 | catGRANULE 2.0 ROBOT | FuzDrop | PICNIC |
|---|---|---|---|---|
| 1 | 0,882 | 0,765 | 0,786 | 0,743 |
| 2 | 0,647 | 0,588 | 0,692 | 1,000 |
| 3 | 0,824 | 0,706 | 0,733 | 0,706 |
| 4 | 0,765 | 0,588 | 0,692 | 0,882 |
| 5 | 0,941 | 0,765 | 0,857 | 0,647 |
| 6 | 0,941 | 0,765 | 0,769 | 0,765 |
| 7 | 0,706 | 0,765 | 0,667 | 0,824 |
| 8 | 0,706 | 0,647 | 0,786 | 0,647 |
| 9 | 0,824 | 0,765 | 0,800 | 0,706 |
| 10 | 0,706 | 0,882 | 0,667 | 0,824 |

| (B) | ParSe_V2 | catGRANULE 2.0 ROBOT | FuzDrop | PICNIC |
|---|---|---|---|---|
| 1 | 0,714 | 0,565 | 0,579 | 0,541 |
| 2 | 0,611 | 0,556 | 0,450 | 1,000 |
| 3 | 0,700 | 0,545 | 0,550 | 0,545 |
| 4 | 0,813 | 0,588 | 0,563 | 0,536 |
| 5 | 0,800 | 0,542 | 0,632 | 0,478 |
| 6 | 0,762 | 0,520 | 0,588 | 0,510 |
| 7 | 0,632 | 0,591 | 0,444 | 0,519 |
| 8 | 0,923 | 0,579 | 0,524 | 0,478 |
| 9 | 0,778 | 0,565 | 0,800 | 0,490 |
| 10 | 0,750 | 0,556 | 0,588 | 0,519 |

| (C) | ParSe_V2 | catGRANULE 2.0 ROBOT | FuzDrop | PICNIC |
|---|---|---|---|---|
| 1 | 0,789 | 0,650 | 0,667 | 0,626 |
| 2 | 0,629 | 0,571 | 0,545 | 1,000 |
| 3 | 0,757 | 0,615 | 0,629 | 0,615 |
| 4 | 0,788 | 0,588 | 0,621 | 0,667 |
| 5 | 0,865 | 0,634 | 0,727 | 0,550 |
| 6 | 0,842 | 0,619 | 0,667 | 0,612 |
| 7 | 0,667 | 0,667 | 0,533 | 0,636 |
| 8 | 0,800 | 0,611 | 0,629 | 0,550 |
| 9 | 0,800 | 0,650 | 0,800 | 0,578 |
| 10 | 0,727 | 0,682 | 0,625 | 0,636 |

**Supplementary Figure 1:** Comparison of the amino acid distribution of sequences without LLPS region against LLPS-driving regions (detected for a window of 50 residues) (Mann-Whitney U p-value ≥ 0.05: N.S., p-value < 0.05: *, p-value < 0.01: **, p-value < 0.001: *** and p-value < 0.0001: ****):

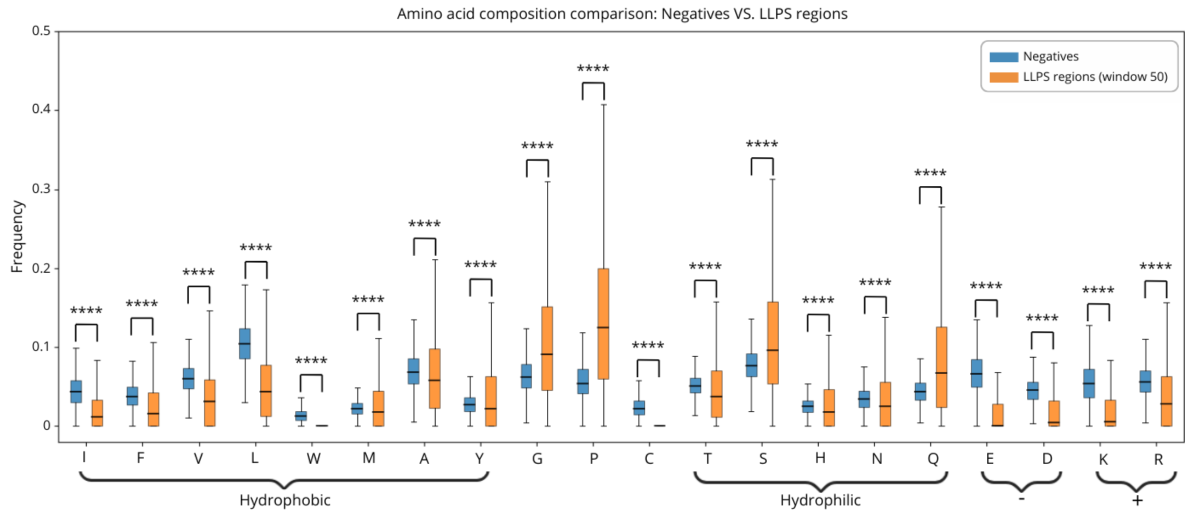


**Supplementary Figure 2:** Comparison of the amino acid distribution of IDRs (IUPred3 score > 0.3) against LLPS regions (window of 50) (Mann-Whitney U p-value ≥ 0.05: N.S., p-value < 0.05: *, p-value < 0.01: **, p-value < 0.001: *** and p-value < 0.0001: ****):

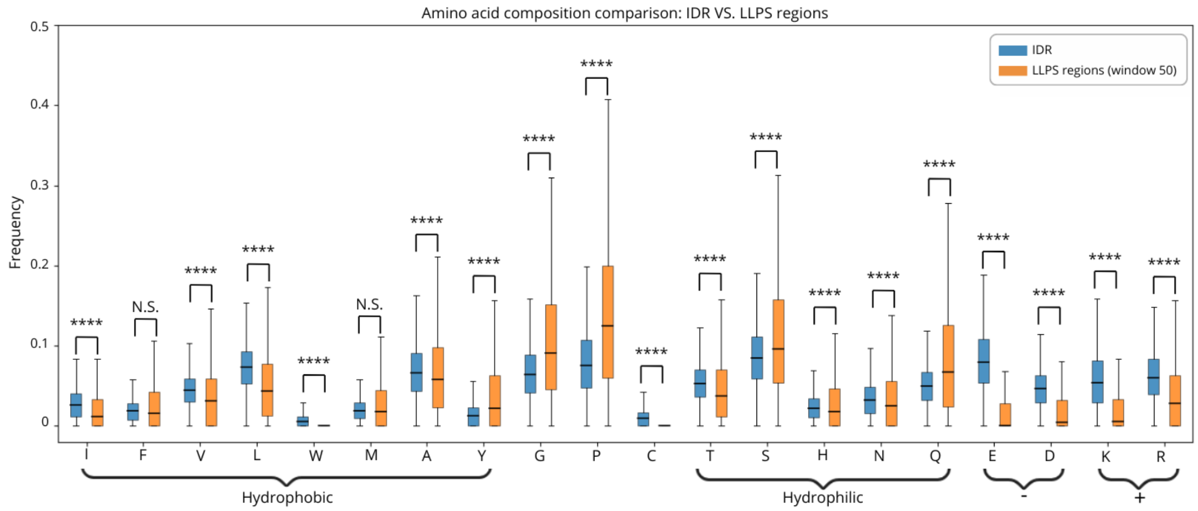
